# A highly parallel, automated platform enabling individual or sequential ChIP of histone marks and transcription factors

**DOI:** 10.1101/728634

**Authors:** Riccardo Dainese, Vincent Gardeux, Gerard Llimos, Daniel Alpern, Jia Yuan Jiang, Antonio Carlos Alves Meireles-Filho, Bart Deplancke

## Abstract

Despite its popularity, chromatin immunoprecipitation followed by sequencing (ChIP-seq) remains a tedious (>2d), manually intensive, low-sensitivity and low-throughput approach. Here, we combine principles of microengineering, surface chemistry and molecular biology to address the major limitations of standard ChIP-seq. The resulting approach, FloChIP, automates and miniaturizes ChIP in a beadless fashion while facilitating the downstream library preparation process through on-chip chromatin tagmentation. FloChIP is fast (<2h), has a wide dynamic range (from 10^6^ to 500 cells), is high-throughput (up to 64 parallel, antibody- or sample-multiplexed experiments) and is compatible with both histone mark and transcription factor ChIP. In addition, FloChIP’s interconnected design allows for straightforward chromatin re-immunoprecipitation, thus constituting the first example of a microfluidic sequential ChIP-seq system. Finally, we demonstrate FloChIP’s high-throughput capacity by performing ChIP-seq of the transcription factor MEF2A in 32 distinct human lymphoblastoid cell lines, providing novel insights into the main factors driving collaborative DNA binding of MEF2A and into its role in B-cell-specific gene regulation. Together, our results validate FloChIP as a flexible and reproducible automated solution for individual or sequential ChIP-seq.

## Main

The genome-wide distribution and dynamics of protein-DNA interactions constitute a fundamental aspect of gene regulation. Chromatin immunoprecipitation followed by next generation sequencing (ChIP-seq)(1) has become the most widespread technique for mapping protein-DNA interactions genome-wide. ChIP-seq has been successfully applied to dozens of transcription factors (TFs), histone modifications, chromatin modifying complexes, and other chromatin-associated proteins in humans and other model organisms(2). The ENCODE and modENCODE consortia alone have already performed more than 8,000 ChIP-seq experiments, which have greatly enhanced our collective understanding of how gene regulatory processes are orchestrated in humans as well as several model organisms(3). In addition, ChIP-seq proved to be essential to acquire new insights into genomic organization(4–6) and into the mechanisms underlying genomic variation-driven phenotypic diversity and disease susceptibility(5, 7, 8). More specifically, this assay proved crucial in determining the DNA binding properties of hundreds of TFs(9). Nevertheless, in comparison to other widespread NGS-based methods – e.g. RNA-seq(10) and ATAC-seq(11) – ChIP-seq lags behind in some key metrics, i.e. throughput, sensitivity, and automation, which hinders its wider adoption and reproducibility. For example, while RNA-seq can now be regularly performed on hundreds or thousands of single cells using readily available workflows(12, 13), ChIP-seq has largely remained labor intensive and limited to few samples per run, each composed of millions of cells. Moreover, while a typical pre-amplification RNA-seq workflow consists of only three steps – i.e. cell lysis, RNA capturing and reverse transcription – ChIP-seq typically involves several pre-amplification steps (crosslinking, lysis, fragmentation, immunoprecipitation, end-repair and adapter ligation). Finally, any given RNA transcript is present in each cell in numerous copies, which increases the likelihood of its capture and detection, whereas, on the other hand, each locus-specific protein-DNA contact occurs a maximum of two times in a diploid cell. The combination of these idiosyncratic differences, together with the lack of enabling solutions, has thus far prevented the ChIP-seq technology, as opposed to other NGS-based methods, to reach its full potential in terms of adoption, utility, and biomedical relevance.

In addition to the standard ChIP protocol, a modification of its workflow involving sequential chromatin immunoprecipitation (sequential ChIP) has been developed to infer the genomic co-occurrence of two distinct (protein) targets. In principle, sequential ChIP consists of performing ChIP twice on the same input chromatin, which leads to a multiplication of the inefficiencies mentioned above. Therefore, not only does sequential ChIP show the same limitations as regular ChIP-seq, but these also come in an augmented form due to its consecutive nature. As a result, few studies have so far performed sequential ChIP followed by next generation sequencing(14, 15) (sequential ChIP-seq) and most of the available studies have so far relied on qPCR to validate putative bivalent regions(16–18) (sequential ChIP-qPCR).

In recent years, several attempts have been made to alleviate some of the limitations of the ChIP-seq and sequential ChIP approaches. Gasper et al.(19) and Aldridge et al.(20) addressed the issue of automation by implementing the manual steps of a conventional ChIP-seq workflow on robotic liquid handling systems. However, in these examples, automation was balanced with sensitivity, since these workflows still require tens of millions of cells per experiment. van Galen et al.(21) and Chabbert et al.(22) addressed the issue of throughput by barcoding and pooling chromatin samples before immunoprecipitation (IP). Although van Galen and colleagues proved that their approach led to higher sensitivity (500 cells per ChIP), neither approach is automated and both are so far limited to the detection of histone marks. Ma et al.(23) and Rotem et al.(24) addressed the limit of sensitivity with two different microfluidic-based strategies. Ma et al. focused on improving the efficiency of the IP step by confining it within microfluidic channels. Although they showed good IP efficiency down to as few as 30 cells, their approach requires impractical antibody-oligo conjugates, is not automated and was not shown to work for TFs. On the other hand, Rotem et al. achieved the remarkable feat of performing ChIP-seq in a single cell by integrating the concept of chromatin barcoding and pooling into a single droplet-based microfluidic chip. However, even though the barcoding step has indeed single cell resolution, the most critical step – i.e. the IP step – is performed manually on 100 cells. As a result, their approach - also shown to work only for histone marks - yielded sparse single cell data and thousands of assays are still required to identify specific cell subpopulation signatures. In a notable effort to simplify the sequential ChIP workflow, Weiner et al.(15) complemented the IP steps with sequential chromatin barcoding, thus achieving a high degree of multiplexing. However, their approach increases the number of experimental steps which makes it significantly more labor-intensive given that the workflow is not automated. Recently, orthogonal approaches have emerged such as CUT&RUN(25) and CUT&Tag(26) which are capable of profiling chromatin in a one-tube format and down to single cells. Rather than relying on solid state separation of protein/DNA complexes, these approaches exploit fusion proteins (protein A-Mnase and protein A-Tn5, respectively) to selectively digest or tagment the genomic DNA in the proximity of chromatin-bound antibodies. Such alternative strategies hold great potential as sensitive and streamlined techniques for genomic profiling of protein/DNA complexes but, as opposed to ChIP-seq, their wide applicability still needs to be proven. Moreover, CUT&RUN and CUT&Tag are unable to probe bivalency, which constitutes a fundamental limit of both techniques. In sum, previous valuable attempts at improving the technology have only addressed a subset of the major ChIP-seq limitations.

In this work, we aimed to address all major limitations of current ChIP-seq and sequential ChIP solutions (throughput, sensitivity and automation), by developing a microfluidic strategy that we named FloChIP. We show that high quality and parallel / multiplexed ChIP-seq for histone marks (down to 500 cells) and TFs (100’000 cells) is achieved in less than two hours through a combination of microvalves, micropillars, flexible surface chemistry and on-chip chromatin tagmentation. Moreover, by designing an interconnected and modular device, FloChIP enables straightforward re-IP of eluted chromatin, effectively establishing a half-day sequential ChIP pipeline. Finally, we demonstrate the high-throughput capabilities of our system by performing ChIP-seq of the TF MEF2A using chromatin derived from 32 lymphoblastoid cell lines (LCLs). Our data highlight the main drivers of MEF2A collaborative DNA binding and provide new insights into MEF2A’s role in the regulatory network underlying lymphoblastoid proliferation.

## Results

### FloChIP is engineered for automated, bead-less and miniaturized ChIP-seq

Conventional ChIP-seq requires hours of manual work, a wide range of consumables provided by a variety of suppliers and has limited sensitivity. The main rationale behind FloChIP’s design was the development of an efficient solution that would address these drawbacks in a convenient and compact manner. The two core elements of FloChIP’s technology are the assembly of a multilayered stack of biomolecules, enabling versatility in antibody pull-down (**Fig. 1a**) and an engineered pattern of high surface-to-volume micropillars for efficient chromatin capture and washing (**Fig. 1b**). FloChIP’s surface chemistry is based on strong although non-covalent molecular interactions and leads to the immobilization of an antibody of choice prior to IP. The first layer is obtained by flowing on-chip a concentrated solution of biotinylated-BSA, which passively adsorbs to the hydrophobic walls of the microfluidic device. This layer has both an insulating role, preventing non-specific adsorption of chromatin to the chip walls, and a docking role for the next layer, which is obtained by flowing on-chip a solution containing neutravidin that strongly binds to the biotin groups of the first layer. The third layer is formed by flowing a solution of biotinylated-protein A/G, which becomes firmly immobilized by the unsaturated binding sites of the neutravidin layer. Protein A/G is a recombinant protein used in a variety of immunoassays due to its ability to strongly bind to a large number of different antibodies. This ability is retained by FloChIP’s surface functionalization which thus constitutes a general substrate for antibody pull-down.

**Fig. 1.**
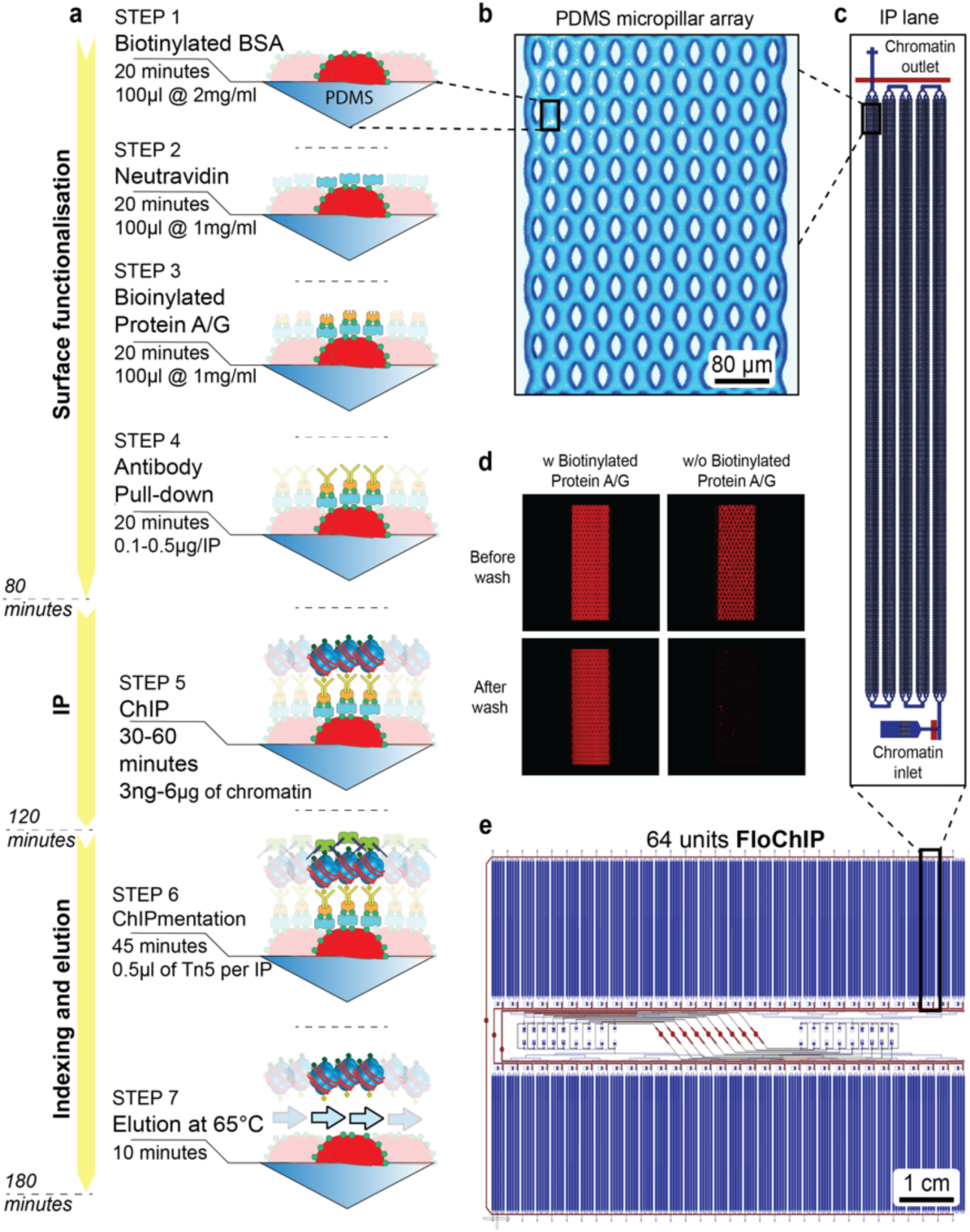
FloChIP’s architecture for miniaturized ChIP-seq. (**a**) FloChIP’s processing phases in descending chronological order. In the first “surface functionalization phase” (∼80 minutes), the inner walls are functionalised by sequentially introducing chemical species that firmly interact with both the previous and following layers of functionalization. Also in chronological order, these species are biotin-BSA, neutravidin, biotin-Protein A/G and antibody. Following functionalization, the IP takes place by flowing sonicated chromatin on-chip in a total time of 30-60 minutes, depending on the chromatin volume that is introduced. Subsequently, the antibody-bound chromatin is tagmented directly on-chip in order to introduce Illumina-compatible adapters. Finally, the tagmented chromatin is eluted off-chip using an SDS-containing buffer and high temperature. (**b**) Top-view microscopy picture of a portion containing numerous micro-pillars. Each portion is itself repeated several times along the length of one IP lane. (**c**) Top-view schematic of one IP lane. Each IP lane can be repeated *n* times across a FloChIP device. Flow channels are in blue and control channels in red. (**d**) Fluorescence micrographs showing the requirement for protein A/G in the correct formation of FloChIP’s functionalization. (**e**) Top-view schematic of the high-throughput 64-unit FloChIP device. Flow channels are in blue and control channels in red.

Another critical feature of FloChIP’s workflow is the microfluidic tagmentation of immunoprecipitated chromatin. In a previous study, tagmentation of bead-bound chromatin (ChIPmentation) was shown to generally increase cost-effectiveness and sensitivity of the ChIP-seq workflow(27). Here, we built upon this concept and adapted it to obtain the first example of ChIPmentation performed directly on chromatin bound to the inner walls of a microfluidic device (**Fig. 1a**). Briefly, this is achieved by flowing a Tn5 solution into the device while heating the chip surface to 37°C, allowing direct on-chip indexing of chromatin-bound DNA. Importantly, microfluidic ChIPmentation streamlines the downstream library preparation workflow and reduces hands-on time (see also below).

For the successful initiation of the multilayered surface functionalization, the only substrate requirement is the hydrophobic surface of the device polymer. Therefore, to maximize the surface-to-volume ratio of our devices, we designed an array of micropillars (**Fig. 1b**), which repeats multiple times across each IP-lane (**Fig. 1c**). With the goal of visually validating the successful assembly of our multilayered on-chip chemistry and to confirm that every layer is essential to this end, we first sought to IP chromatin derived from a HeLa H2B-mCherry cell line using an anti-H2B antibody. The resulting fluorescence micrographs confirmed that each layer of the molecular species is necessary for successful IP of cellular chromatin (**Fig. 1d, Supp. Fig. 1a)**.

The IP-lane (**Fig. 1c**) is the fundamental unit of the FloChIP architecture and it can itself be repeated *n* times, where *n* is the desired throughput of the device. For our initial tests, we used an 8-lane FloChIP device (**Supp. Fig. 1b-c**), and we later adopted a 64-unit device for higher-throughput experiments (**Fig. 1e, Supp. Fig. 1d**). To gain accurate flow control, automation and multiplexing, a network of soft microvalves was added to the design: different multiplexing modes can be achieved with the same microfluidic architecture by actuating distinct sets of valves. For instance, we named “FloChIP mode 1 - sample multiplex”, the option of coating all IP lanes of the device with one antibody and introducing different samples from dedicated individual inlets (**Fig. 2a**). Alternatively, “FloChIP mode 2 - antibody multiplex” provides the option of coating each IP lane with a different antibody and of distributing one sample equally across the whole device (thus probing distinct antibodies in parallel using one sample) (**Fig. 2b**).

**Fig. 2.**
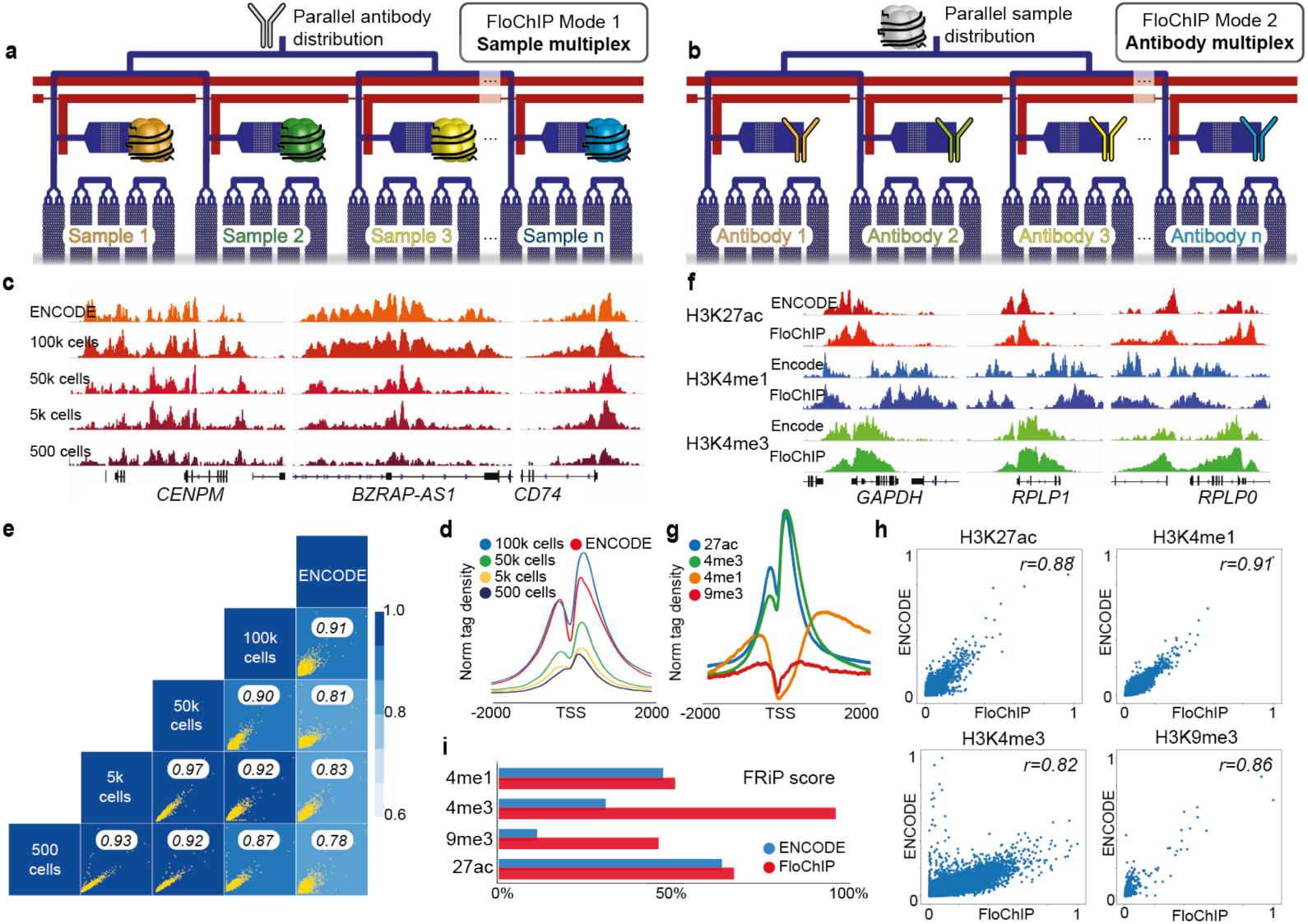
FloChIP-based derivation of chromatin landscapes. (a) Schematic depiction of FloChIP’s *mode 1*: sample multiplex. One antibody solution is introduced through the common inlet and distributed equally across all IP lanes. During IP, each IP lane is loaded separately by introducing different samples through the individual inlets. (**b**) Schematic depiction of FloChIP’s *mode 2*: antibody multiplex. Each IP lane is functionalized separately by introducing different antibodies through the individual inlets. During IP, one sample is introduced through the common inlet and distributed equally across all IP lanes. (**c**) H3K27ac profiles at three different genomic loci obtained by FloChIP with decreasing cell numbers (100k to 500 cells). For comparison, ENCODE data generated by conventional ChIP-seq are also shown. (**d**) Normalized read density meta-profiles around transcription start sites for samples of decreasing cell numbers and ENCODE. (**e**) Genome-wide correlation between pairs of samples with decreasing cell numbers and ENCODE. (**f**) Signal tracks for H3K27ac, H3K4me1 and H3K4me3 profiles obtained by FloChIP with 100’000 cells are shown at three different genomic loci. (**g**) Normalized read density meta-profiles around transcription start sites for H3K4me3, H3K27ac and H3K4me1. For comparison, ENCODE data generated by conventional ChIP-seq are also shown. (**h**) Genome-wide correlation plots between FloChIP (x axis) and ENCODE (y axis) data for all targets tested, i.e. H3K27ac, H3K4me1, H3K9me3, H3K27me3 and H3Kk4me3. (**i**) Comparison in terms of fraction of reads in peaks (FRiP) between FloChIP and ENCODE for histone mark samples.

### FloChIP reliably reproduces ENCODE data across a wide range of input cells

To evaluate the performance of our FloChIP strategy, we first set out to empirically estimate FloChIP’s dynamic range. To this end, we performed FloChIP in sample multiplex mode, by functionalizing the whole chip with an anti-H3K27ac antibody and immunoprecipitating different chromatin dilutions (equivalent to 1 million and 500 cells, respectively). Despite the observed differences in recovered DNA (**Supp. Fig. 2a**), we obtained high and stable fold enrichment results across the whole series of dilutions tested (**Supp. Fig. 2b**). To obtain a genome-wide perspective on its dynamic range, we sequenced FloChIP’s libraries for chromatin samples obtained from 100’000, 50’000, 5’000 and 500 cells. After sequencing, the rate of uniquely mapped reads remained high for all samples (**Supp. Fig. 2c**), while the fraction of reads falling into peaks (FRiP score) decreased with decreasing input amounts – from over 60% for the largest sample, to just above 10% for the smallest (**Supp. Fig. 2d**). Nevertheless, both locus-specific inspection and genome-wide analysis of the obtained libraries revealed reproducible profiles (**Fig. 2c**) and characteristic accumulation of reads into regions in proximity of transcription start sites (TSS, **Fig. 2d**). Moreover, the pairwise correlation of reads in peaks demonstrated the high accuracy of our approach since we uncovered a high correlation between all library pairs (between *R*^*2*^*= 0.78* and *R*^*2*^*= 0.97*). This included ENCODE-FloChIP pairs among which the highest correlation was obtained for the 100’000 cell sample, i.e. *R*^*2*^*= 0.91* (**Fig. 2e**). In addition, to evaluate individual chip-to-chip variation, we analyzed the correlation between two libraries obtained from 100’000 cells in identical conditions but derived from different FloChIP devices. Again, we obtained high correlation (*R*^*2*^*= 0.98*, **Supp. Fig. 2e**), indicating that our system is robust to batch variability.

Next, we set out to evaluate the reproducibility of our approach with other genomic targets. To this end, by using FloChIP’s mode 2: “antibody multiplex”, we performed ChIP-seq of four histone marks in parallel (H3K27ac, H3K4me3, H3K4me1 and H3K9me3) using the same sample, going from chromatin to sequencing-ready libraries, in one day. Following high qPCR enrichment (**Supp. Fig. 2f** and sequencing, we found that the obtained signal tracks closely resemble those of ENCODE (**Fig. 2f**). In addition, to evaluate FloChIP’s performance with greater resolution, we determined the extent of genome-wide distribution of reads around the TSS (**Fig. 2g**) and the correlation between FloChIP and ENCODE datasets. Comparison of signal intensities between the respective datasets confirmed an overall high read density correlation in peaks (H3K4me3: *r = 0.82*, H3K27ac: *r = 0.88*, H3K4me1: *r = 0.91*, H3K9me3: *r = 0.86*; **Fig. 2h**). Moreover, comparison in terms of the FRiP score showed that, despite the ChIP cell input for ENCODE being two orders of magnitude greater than that of FloChIP, our technology consistently yields highly enriched libraries, with FRiP scores between 1.1x and 4.1x higher for FloChIP compared to ENCODE (**Fig. 2i**). These data show that FloChIP can be used to robustly generate chromatin landscapes for histone marks on the same sample in a parallelized manner and over a wide input range.

### FloChIP “sequential IP” mode provides genome-wide information on bivalent regulatory regions

Conventional ChIP-seq provides information on the genome-wide localization of one specific protein or histone modification at a time. However, DNA regulatory elements tend to be characterized by much more complex chromatin states that involve multiple histone marks and collaborating TFs(5, 16, 28). For instance, it has been shown that promoters showing both repressive (H3K27me3) and activating (H3K4me3) marks are a characteristic feature in embryonic stem (ES) cells(16, 29). This class of promoters was originally named “bivalent”(16, 29) and is strongly associated with key spatially regulated developmental genes(30). To obtain direct information on the genomic location of bivalent promoters, a variant of the standard ChIP protocol called sequential ChIP was developed(16). Despite the advantage of sequential ChIP over standard ChIP in discerning true bivalency, its manual involvement and laboriousness have thus far prevented widespread usage. To address the technical limitations of the current sequential ChIP workflow, we exploited FloChIP’s intrinsic modularity, highly efficient IP and multiplexing features to generate, to our knowledge, the first example of an automated and miniaturized sequential ChIP solution (**Fig. 3a**). Briefly, FloChIP’s “sequential IP” consists of two consecutive IPs taking place in two adjacent IP-lanes. The chromatin immobilized and washed in the first IP-lane is resuspended by means of a peptide elution strategy(14) and then transferred on-chip to a neighboring IP-lane where the second IP is carried out (**Fig. 3a**), followed by salt washes and tagmentation.

**Fig. 3.**
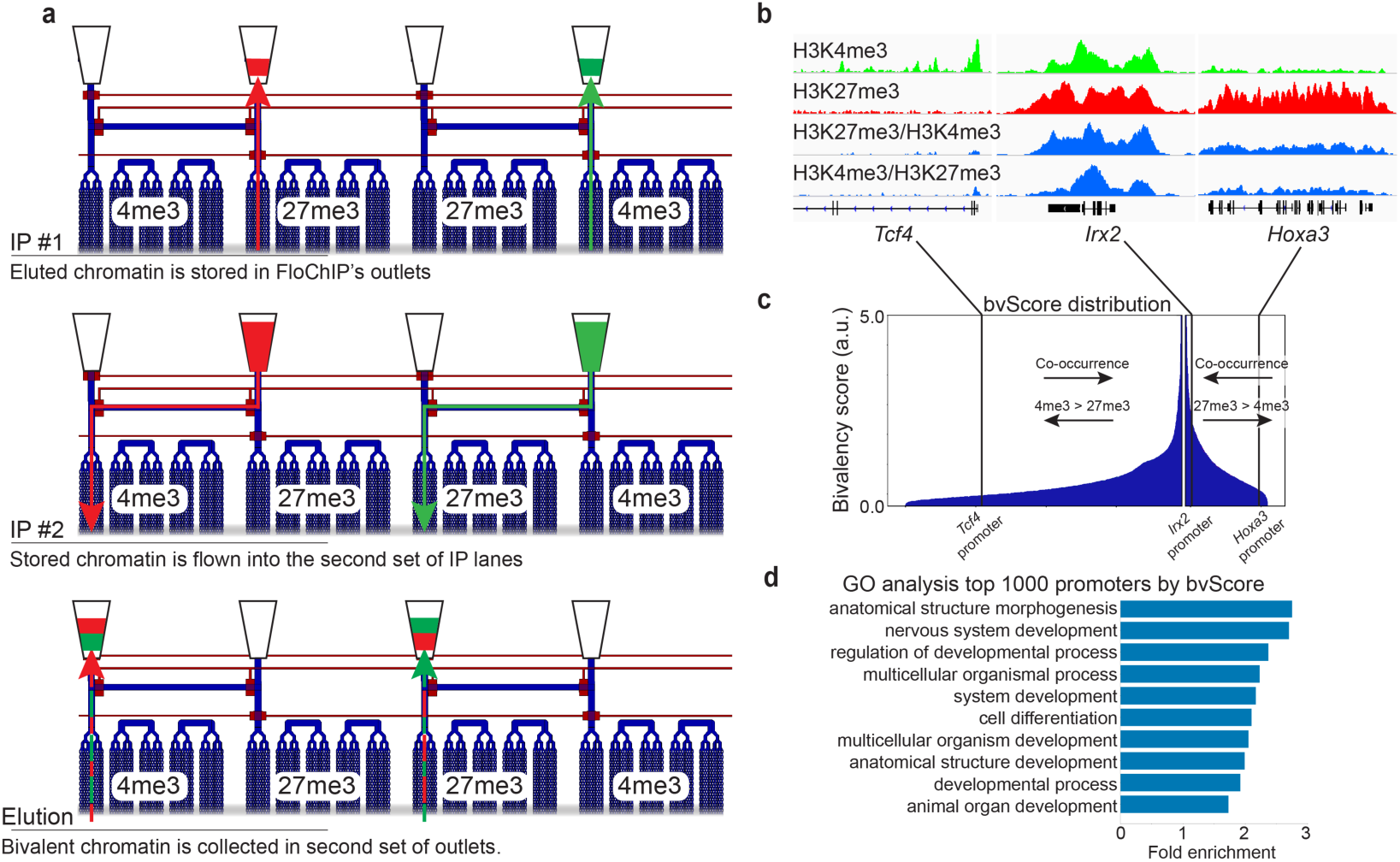
FloChIP’s “sequential IP” mode for the study of factor co-occupancies including bivalent chromatin. (**a**) FloChIP’s sequential IP steps in descending chronological order as applied for H3K4me3-H3K27me3 co-occupancy. Chromatin that is derived from the first IP is collected into off-chip reservoirs connected to the device. Following collection, the control channels are actuated to isolate the first IP lane from the chromatin, while opening the path to the second IP lane. At this point, the chromatin flows into the second pre-functionalised IP lane. Finally, the bivalent chromatin is eluted again in off-chip reservoirs. (**b**) Signal tracks for the two individual IP libraries (H3K4me3 and H3K27me3) as well as the corresponding sequential IP samples (H3K27me3/H3K4me3 and H3K4me3/ H3K27me3) at three different genomic loci. (**c**) Bivalency score values for high-CpG promoters. The color codes reflect the relative abundance of the two individual marks for each considered promoter. (**d**) Gene Ontology enrichment analysis for the first one thousand promoters with the highest bivalency score.

We validated this approach by focusing on bivalent chromatin in embryonic development, given its well-studied role in this context. Specifically, we acquired genome-wide direct co-occupancy profiles for H3K27me3 and H3K4me3 in mouse embryonic stem cells (mESCs) in both IP directions – i.e. H3K27me3 first followed by H3K4me3 (H3K27me3/H3K4me3) and vice versa. As mentioned above, H3K4me3 and H3K27me3 bivalency has been originally attributed to promoters of developmental genes, leading to the hypothesis that a bivalent state maintains genes in a poised state(16). In previous genome-wide sequential ChIP studies, promoters were assigned to bivalency classes, e.g. pseudo bivalency, partial bivalency and full bivalency(14). However, to consider a more plausible continuous distribution of bivalency across promoters, we computed for each promoter a “bivalency score” (bvScore, details in the **Methods** section) instead of assigning them into discrete classes. To evaluate the performance of FloChIP’s “sequential IP” mode, we focused on three distinct regions that have been previously used as proof-of-concept models by Bernstein and colleagues(16), who illustrated the methylation status difference of these regions by using ChIP-qPCR and sequential ChIP-qPCR. With this approach, they were able to distinguish regions displaying only H3K4me3 (e.g. *Tcf4* TSS) and only H3K27me3 (e.g. upstream of *Hoxa3*) versus those displaying true bivalency (e.g. *Irx2* TSS). FloChIP-based genomic profiles (**Fig. 3b, Supp. Fig. 3a**) and bvScore distributions (**Fig. 3c**) validated these previous findings(16). We observed that, as expected, the *Tcf4* promoter shows high H3K4me3 but low H3K27me3 enrichment, and thus low bivalency (*bvScore=0.83)*. On the other hand, *Hoxa3* was mainly marked by H3K27me3, with low H3K4me3 enrichment and consequently low bivalency signal (*bvScore=0.44)*. Finally, the TSS of *Irx2* showed high bivalency (*bvScore=3.34*), with all four genomic tracks, two individual and two sequential FloChIPs, showing high coverage. In addition to considering specific loci, we also validated our data on a genome-wide scale by achieving high correlation with the results obtained by Weiner et. al.(15) using their Co-ChIP system (**Supp. Fig. 3b**). Finally, as another independent validation of our analysis, we performed Gene Ontology enrichment on the first one thousand promoters with the highest bivalency score. As expected, we found that these promoters are highly enriched in genes involved in a number of developmental processes, from anatomical structure development to neurogenesis (**Fig. 3d**). Taken together, our data indicates that FloChIP’s “sequential IP” mode constitutes a miniaturized, low-input (100’000 cells) and rapid (between 5-6 hours) sequential ChIP-seq workflow for the analysis of co-occurring binding events genome wide.

### FloChIP is capable of ChIPing TFs in “high-throughput” mode

As already mentioned above, previous attempts at improving either the sensitivity or multiplexing ability of ChIP-seq experiments were successful but only so far in the context of histone modifications (21–24). The reason for this is that performing ChIP on TFs poses additional challenges compared to ChIP on histone marks. These include the fact that i) TF/DNA interactions are less abundant and weaker than histone mark/DNA interactions and ii) antibodies for TFs normally show lower affinity for their epitopes compared to histone mark antibodies. These challenges translate into the need for more abundant sample inputs and longer incubation times. Indeed, during FloChIP optimization, we also experienced these challenges, rendering FloChIP’s indirect method – i.e. with 2-4 hours antibody/chromatin pre-incubation in tubes – to be the only robust way to obtain high quality TF ChIP results (data not shown). Nevertheless, flowing the pre-incubated antibody/chromatin mixture on-chip, we succeeded in performing IP on the Myocyte Enhancer Factor 2A (MEF2A) TF involving only 100’000 cells, proving for the first time the feasibility of miniaturized and automated TF ChIP (**Fig. 4**).

**Fig. 4.**
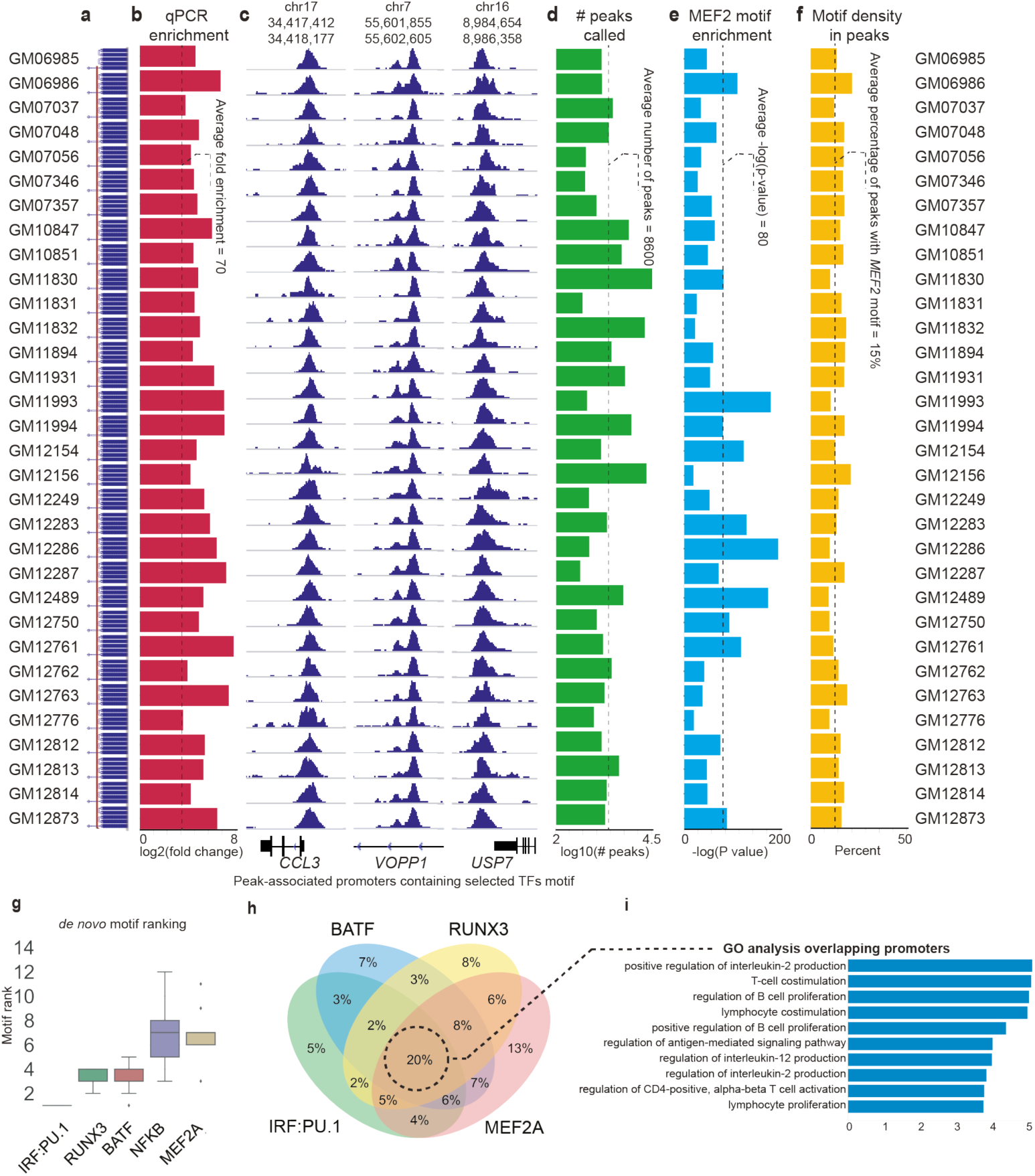
FloChIP’s TF IP high-throughput mode. (**a**) List of the 32 cell lines used in this study. (**b**) qPCR enrichment for each library at the *VOPP1* locus. The average fold enrichment across all libraries is 70, represented as a dotted line. (**c**) Signal tracks are reported for each library for three different genomic loci. (**d**) Number of peaks called for each library (8600 peaks on average, represented by a dotted line). (**e**) MEF2A motif enrichment for each library (the median p-value is 1e-10, represented as a dotted line). (**f**) Percent of called peaks containing the MEF2A motif (average is 15%, represented as a dotted line). (**g**) Results of *de novo* motif search and ranking based on the respective enrichment of each detected motif across libraries. (**h**) Venn diagram showing the percentages of MEF2A-bound promoters featuring no or occurrence of other detected motifs. (**i**) Gene ontology enrichment analysis of promoters containing MEF2A, BATF, RUNX3 and IRF:PU.1 motifs.

While demonstrating the applicability of our approach on TFs, we also set out to demonstrate the high-throughput capabilities of our device. To this end, and by using half of the 64 IP lanes of the FloChIP device, we performed MEF2A ChIP-seq on chromatin derived from distinct LCLs derived from 32 unrelated European individuals whose genomes were sequenced as part of the 1000 Genomes Project(31) (**Fig. 4a**). We specifically targeted this TF given its association with variable chromatin modules that were inferred from histone mark and PU.1 ChIP-seq data from LCLs, as presented in one of our previous studies(4). Before sequencing, we verified the IP quality of each library by qPCR (**Fig. 4b**). Fold change results indicated consistent, high enrichment across the 32 IP lanes (*avg(Fold enrichment)=70*). In addition, we noticed that, during library preparation, the number of amplification cycles that was required to obtain sufficient DNA amounts for NGS sequencing was the same for all 32 samples (17 PCR cycles), reflecting homogeneity in DNA yield. This result was obtained without manually adjusting the volume or concentration of the sample prior to IP. We therefore reasoned that FloChIP itself, when used in saturating conditions, provides the additional advantage of equalizing the amount of recovered DNA, which renders the library preparation process straightforward. After sequencing, we confirmed the accumulation of mapped reads in selected genomic loci (based on MEF2A ENCODE data: upstream *CCL3, VOPP1* and upstream *USP7*; **Fig. 4c**).

Subsequently, we analyzed the quality of our data using a number of genome-wide measures. First, we called peaks for each sample (**Fig. 4d**, *avg(#peaks)=8600*) and observed a general high degree of genome-wide read density correlation for all sample pairs (*avg(cor)=0.84*) (**Supp. Fig. 4a**). Surprisingly, despite detecting strong enrichment of the MEF2-like consensus motif in each individual peak set (**Fig. 4e**, *median(pval)=1e-10*), examining the presence of motifs in peaks revealed that only a small portion of peaks contained the MEF2A motif (**Fig. 4f**, *avg(motif density in peaks)=15%*). Therefore, we set out to explore alternative drivers of the observed MEF2A binding. To this end, we performed *de novo* motif discovery on each individual peak file and ranked the obtained motifs according to their statistical significance. Interestingly, the MEF2A motif emerged as only the fifth most enriched one, preceded by the motifs for BATF, RUNX3, NFKB and the IRF:PU.1 dimer (**Fig. 4g**). This observation is consistent with previous motif-based analyses in LCLs (4, 32) and suggests that MEF2A DNA binding could be largely motif-independent and driven by the interaction with other factors(33). To note, a previous study aimed at deciphering the Epstein–Barr-Virus-based mechanism of B-cell/lymphoblastoid conversion had also found the same motifs to be strongly enriched in their EBNA3C ChIP-seq dataset(32). Thus, we hypothesized that, in LCLs, MEF2A may play a role in the gene regulatory network underlying B-cell proliferation. To test this hypothesis, we first split the observed peaks in unique and overlapping sets based on the presence of one or more of the MEF2A motifs and the three other motifs with the highest enrichment, i.e. IRF:PU.1, RUNX3 and BATF (**Fig. 4h**). We found that the largest set of the resulting Venn diagram consists of peaks containing all four motifs (19% of all peaks) and is enriched for ontology terms linked to immune cell regulation and proliferation. (**Fig. 4i, Supp. Fig. 4b**). In comparison, the second largest set (13%) involved peaks featuring exclusively the MEF2A motif and did not show insightful ontology enrichment. These findings indicate that MEF2A may play a role in the gene regulatory network underlying lymphoplastoid cell proliferation and that it does so specifically as part of a larger complex of collaborating TFs.

Next, we examined the impact of genetic variation on MEF2A binding. To achieve this, we exploited our large MEF2A ChIP-seq dataset by considering allele-specific binding events (ASBs, see Methods) across all 32 samples and found that only a very small fraction (0.7%) of ASBs could be explained by MEF2A motif variation (**Fig. 5a, Supp. Fig. 5a-d**). Indeed, only 5 out of 751 ASBs were significantly disrupting or creating a MEF2 motif (considering the entire MEF2 family) at FDR < 5% and were concordant in terms of ASB allele effect. Similarly, we found 3 ASBs for *BATF*, 19 for *RUNX*, and 2 for *IRF* (∼4% of all ASBs). In total, we tested 401 mononucleotide human core TF motifs (from HOCOMOCO v11(34)), revealing only 223 ASBs that were significantly disrupting at least one motif at FDR5% (∼30%). This result is consistent with our observation that few MEF2A DNA binding events are dependent on its own motif and also aligns with the notion that the majority of variable TF DNA binding events is driven by motif-independent mechanisms(5, 33).

**Fig. 5.**
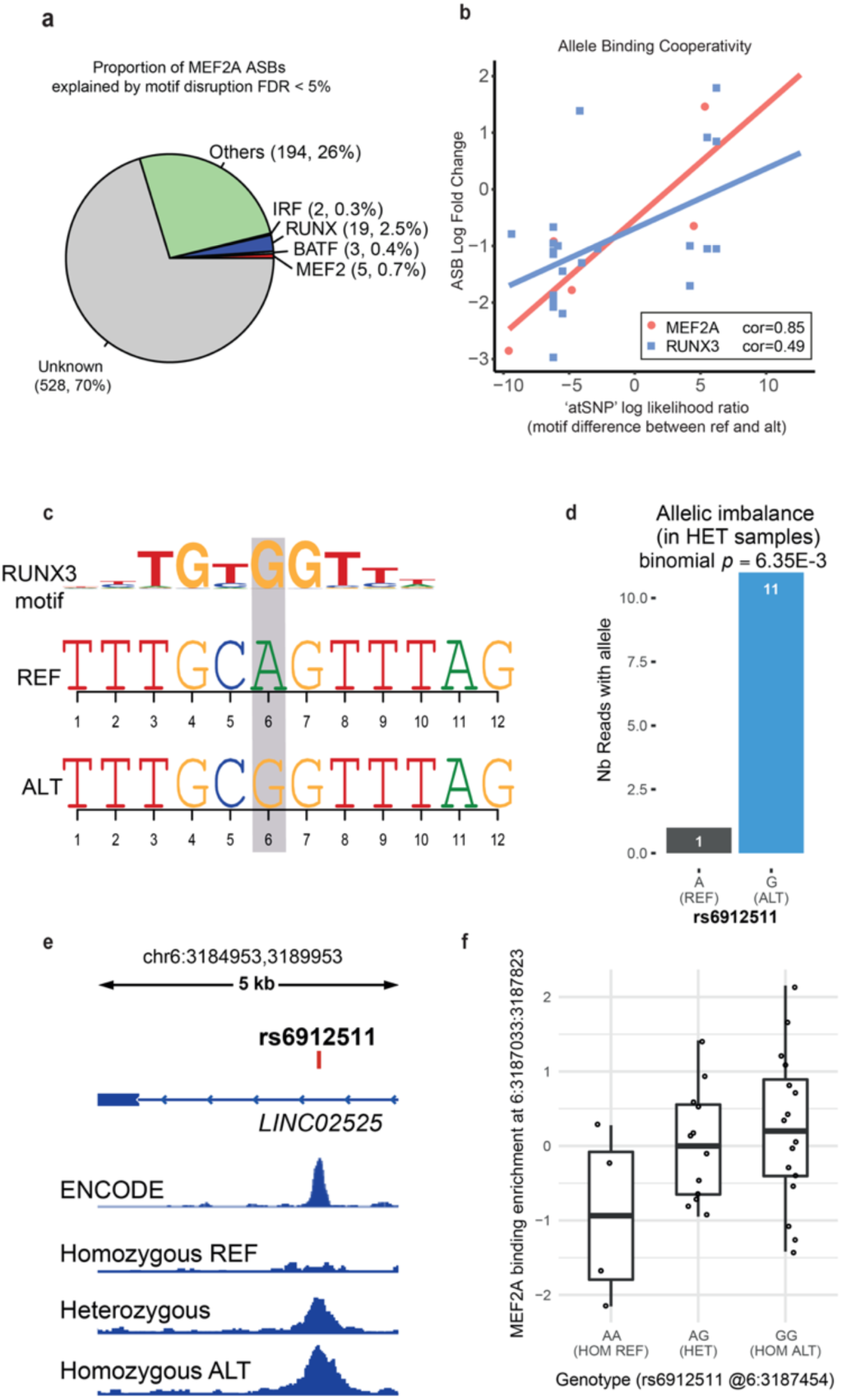
Genetic variation-based analyses reveal RUNX3 to be an important mediator of MEF2A DNA binding. (**a**) Proportion of the 751 identified ASBs that can be explained by motif disruption (at FDR5%). RUNX, IRF and BATF sum the results of all the TF members in their family (because the respective motifs are very close). Only ASBs that are concordant with motif disruption or creation are counted. (**b**) Allelic Binding Cooperativity (ABC) test for inferring motif disruption of other TFs that are modulating MEF2A DNA binding. The x-axis represents the motif score/quality difference (for MEF2A in red, and RUNX3 in blue), while the y-axis represents the fold change between the number of reads mapping to the ref vs the alt allele, summed over all heterozygous samples. This was computed for all 5 identified MEF2A ASBs, and 22 RUNX3 ASBs (at FDR 5%). (**c**) rs6912511 significantly disrupts the RUNX3 motif motif (FDR <5%). (**d**) Allelic Imbalance highlighted for rs6912511 by summing read counts of each allele over all heterozygous samples. A binomial test yielded a p-value of 6.35E-3 which revealed rs6912511 as an ASB. (**e**) IGV tracks showing MEF2A binding enrichment at rs6912511 loci. The coverage tracks are from ENCODE (NA12878 MEF2A) and our 32 samples, stratified by rs6912511 genotype, and merged into three tracks (final bigwigs are RPKM-normalized using deepTools). (**f**) Boxplots showing the effect of rs6912511 on the three different genotypes (AA, AG and GG). MEF2A binding enrichment (y-axis) was computed from the peak in which rs6912511 was found, and normalized using DESEq2, followed by qqnorm functions in R.

Interestingly, and as indicated above, we found that a large portion of motif-affected ASBs are linked to RUNX-family binding sites (**Fig. 5a**). In agreement with this, allelic binding cooperativity analysis showed that variation in RUNX-like motifs was significantly correlated with MEF2A DNA binding differences (nominal p-val < 5%), pointing to cooperative DNA binding mechanisms between these TFs (**Fig. 5b**). Indeed, an Allelic Binding Cooperativity (ABC, see Methods) test revealed that RUNX3 (with 22 SNPs tested) is one of the top significant co-binders with MEF2A (p-value = 0.02). MEF2A and MEF2D motifs also appeared in the top 7 list with p-values of 0.068 and 0.051 respectively. Of note, since the motifs of RUNX factors are very similar to one another, our results cannot specifically distinguish between RUNX1, RUNX2 and RUNX3 (data not shown). Nevertheless, given that only RUNX3 ChIP-seq data exists for LCLs (https://www.encodeproject.org/targets/RUNX3-human/) and that previous evidence also linked RUNX3 to B-cell proliferation pathways(32), we narrowed our analyses to this TF. Interestingly, within the set of RUNX3 ASBs, we detected several cases in which genetic variation from the reference sequence either creates (**Fig. 5c**) or disrupts a RUNX motif (**Supp. Fig. 5e**). Accordingly, RUNX motif creation induced MEF2A DNA binding to the alternate allele (**Fig. 5d-f**), whereas motif disruption led to MEF2A ChIP signal loss (**Supp. Fig. 5f-h**). Taken together, these genetic variation-based results suggest that within the TF complex governing lymphoblastoid cell proliferation, RUNX3 and MEF2A cooperate, with RUNX3 acting as a key driver of MEF2A DNA binding. Together, these results demonstrate the value of FloChIP in catalyzing the relatively straightforward acquisition of TF binding data across many genotypes.

## Discussion

Profiling the interactions between proteins and DNA has both fundamental and biomedical value(2, 35, 36), but continues to constitute an important technological challenge for genomics research(3, 37). ChIP-seq allows the probing of protein-DNA interactions on a genome-wide scale, thus achieving high-throughput in terms of DNA sequence space coverage. However, in terms of experimental output, ChIP-seq largely remains a low-throughput technique. This is mainly due to the long and manually intensive ChIP-seq pipeline which, although widespread, only offers limited automation, reproducibility and sensitivity. Valuable efforts have already been devoted to addressing ChIP-seq’s main limitations but these efforts tended to target only specific issues while ignoring others, as outlined in the introduction (14, 15, 18, 20–24, 27). To more comprehensively address the limitations of standard ChIP-seq, we developed a novel, microfluidic system that we named FloChIP. In this work, we demonstrate that FloChIP enables both rapid single and sequential IP across a wide input range, target flexibility, and experimental scalability.

We first characterized FloChIP’s dynamic range by targeting H3k27ac in samples containing 500 up to 1’000’000 cells. We show high fold enrichment and good genomic coverage for all tested samples, indicating that FloChIP performs robustly across a wide cell number range. In a second step, we tested different antibodies in what we define as the “antibody multiplex mode”. In this configuration, the different IP-lanes of our device are individually functionalized with distinct antibodies with one sample being distributed in parallel to all immunoprecipitation units. The geometric layout of the microchannels thereby ensures uniform distribution of the sample (**Fig. 1e** and **Supp. Fig. 1b**). After observing good correlations with the benchmarking data (ENCODE) for all tested histone mark targets, we conclude that FloChIP can generate chromatin state landscapes with reliability and flexibility.

After establishing this proof of concept on histone marks, we set out to expand the applicability of FloChIP in two directions: sequential ChIP-seq and ChIP-seq on TFs. For the former, we took advantage of the use of microvalves to compartmentalize distinct sections of the microfluidic device in a controllable manner, as reported previously for other applications(38, 39). More specifically, the use of microvalves allows to easily orchestrate the functionalization of adjacent IP lanes with different antibodies, in this case H3K4me3 and H3K27me3, and to transfer eluted chromatin from one IP lane to the next. With this approach, we recapitulated the landmark qPCR(16) and sequencing(15) results on the genomic distribution of H3K4me3/H3K27me3 bivalency in embryonic stem cells. Here, we note that the key advantage of FloChIP lies in turning sequential ChIP assays from a long (>2 days), intensive, and error-prone protocol into a fast (half day) and automated procedure. We therefore believe that FloChIP could catalyze renewed interest in histone mark bivalency(15, 17), whose molecular function and relevance remains poorly understood in part because of the cumbersome nature of the sequential ChIP-seq technique.

Consistent with the general purpose scope of our microfluidic ChIP-seq system, our final goal was to demonstrate the ability of FloChIP to carry out immunoprecipitation on TF targets. Such capacity would constitute important technical progress, since previous microfluidic or multiplexing implementations of ChIP focused exclusively on histone marks(21, 23, 24). To cope with the generally lower affinity of TF antibodies for their targets, we complemented the regular FloChIP procedure with antibody-chromatin pre-incubation and oscillatory sample loading(40). Using a high-throughput FloChIP device, we completed 32 MEF2A immunoprecipitations and on-chip tagmentations in one run and in less than 4h (including 2 hours of pre-incubation). To the best of our knowledge, no other available solution has so far demonstrated the same level of automation and throughput. After sequencing, we used the data to investigate the role of MEF2A in the gene regulatory network underlying lymphoblastoid cell proliferation. By integrating genotypic and molecular phenotypic data, we found that MEF2A DNA binding is in large part controlled by other TFs such as RUNX3, which, as part of a larger complex of collaborating TFs, seem to coordinate B-cell proliferation. Importantly, both sequential ChIP-seq and TF ChIP-seq experiments were carried out on samples of 100’000 cells, therefore constituting a significant improvement in sensitivity for these ChIP-seq variants, which usually require several millions of cells(3).

Finally, we anticipate that FloChIP can still benefit from further optimization. Despite its ability to automate several steps, such as IP, washes and tagmentation, FloChIP demands important preparatory hands-on work, e.g. wiring the device and interfacing it to the control system. Moreover, even though FloChIP streamlines a significant portion of the ChIP-seq workflow, it remains sensitive to the pre-IP protocol steps and to the choice of specific antigen targets. In other words, similar to standard ChIP-seq, high-affinity antibodies and correctly fragmented chromatin will remain essential requirements for the correct functioning of FloChIP. Future efforts will therefore be tailored towards expanding the range of steps performed directly on the device, such as cell lysis and chromatin fragmentation, for instance, which will yield a truly end-to-end solution.

In conclusion, we believe that FloChIP has the potential to empower the community with a practical and reliable immunoprecipitation solution. Its future integration with standardized user-friendly devices will thereby pave the way toward full automation.

## Methods

Detailed info can be found in the Supporting methods section of the SI appendix

### Chromatin preparation

Lymphoblastoid cells were harvested, washed once with PBS and resuspended in 1ml crosslinking buffer. Crosslinking was quenched and cells were then washed twice with ice-cold PBS, pelleted, deprived of the supernatant, snap frozen and stored at −80°C. The frozen cell pellet was resuspended in ice-cold PBS, lysed and resuspended in sonication buffer. Nuclei were sonicated on a covaris E220 machine.

### FloChIP

FloChIP devices were fabricated as previously reported(41) (details in the Supporting methods).

A FloChIP experiment starts with pre-loading the control lines with distilled water and activating all valves. Subsequently, all the reagents required for the surface chemistry are loaded into pipette tips and inserted into the inlets of the microfluidic device. At this stage, all valves are closed and there is no possible cross-talk between any of the reagents above. Immediately after completing the insertion of the tips, the automated protocol is launched by running the respective script. The protocol entails, in sequential order, the following steps: 20 minutes of BSA-biotin (100μl at 2mg/ml), 30 seconds of PBS wash, 20 minutes of Neutravidin (100μl at 1mg/ml), 30 seconds of PBS wash, 20 minutes of biotin-protein A/G (100μl at 2mg/ml) and 30 seconds of PBS wash. The antibodies used in this study are: Abcam antibodies: anti-H3K27ac ab4729, anti-H3K4me3 ab8580, anti-H3K4me1 ab8895, anti-H3K9me3 ab8898, anti-H3K27me3 ab6147, and anti-MEF2A sc-17785. Following antibody loading chromatin samples are loaded on chip by opening and closing the respective microvalves. Following IP, salt washes are performed to eliminate non-specific binding. Subsequently, Tn5 buffer is flown on-chip to tagment the immunoprecipitated chromatin.

Following Tn5 buffer, SDS is loaded on-chip at 65°C for 10 minutes in order to elute the antibody-bound chromatin from the device. The eluate is independently collected from each IP lane into PCR tubes and de-crosslinked at 65°C for 4 hours. Following de-crosslinking, DNA is purified in Qiagen EB buffer using Qiagen MinElute purification kits.

### FloChIP read mapping and processing for histone marks and TFs

Sequencing reads were mapped to the human (hg38) and mouse (mm10) genomes using STAR(42) with default parameters. Uniquely mapped reads were used to call peaks using the HOMER(43) command *findPeaks.pl* with the appropriate flag, i.e. *–histone* for histone marks and *–factor* for transcription factors. FRiP scores were calculated using HOMER’s command *annotatePeaks.pl*, dividing the total number of reads that fall within peaks by the total number of mapped reads. Correlation plots were generated using *annotatePeaks.pl*.

### Allele-Specific Binding

For identifying variants subject to Allele-Specific Binding of MEF2A (ASB), we downloaded the 1000G genotyping data from the <u>EBI server</u> (hg19), removed variants with a minor allele frequency lower than 5%, while restricting our analysis to the 32 samples of interest, which yielded a total of 1,268,985 single nucleotide polymorphisms (SNPs). Then, we applied the *ASEReadCounter* tool from GATK v.4.0.4.0, on each of the 32 samples. These results were then merged by summing read counts for every SNP across heterozygous samples. We filtered out all SNPs with a coverage lower than 10 reads, which yielded 6,330 SNPs. Then we opted to keep only SNPs falling into called peaks, which led to 4,554 SNPs that were further analyzed. On these, we performed a binomial test to assess significant allelic imbalance (nominal p-value 5%), yielding 751 potential ASBs (37 at FDR 5%).

### TF motif disruption analysis

All downstream analyses were performed using R v. 3.5.0. We analyzed all 751 ASBs to check if they were significantly impacting Transcription Factor (TF) motifs using atSNPv.1.0(44) with the *BSgenome.Hsapiens.UCSC.hg19* package as genome library and *SNPlocs.Hsapiens.dbSNP144.GRCh37* package as SNP library. We tested 401 mononucleotide human core TF motifs that were downloaded from <u>HOCOMOCO v11</u>(34).

### Allele Binding Cooperativity

Allele Binding Cooperativity (ABC) was assessed for all 751 ASBs using a linear regression between the motif disruption/creation log likelihood ratio computed by atSNP (see previous Methods section) and the fold change between the ASB ref and alt counts. This analysis was performed for all SNPs significantly disrupting any of the 401 motifs analyzed by atSNP at FDR5%, and allowed to find which ASBs were concordant with motif disruption across several tested SNPs.

## Acknowledgments

This work has been supported by funds from the Swiss National Science Foundation (#31003A_162735 and #CRSII3_147684), by SystemsX.ch Special Opportunity Project 2015/323, and by Institutional support from the Ecole Polytechnique Fédérale de Lausanne (EPFL).

