## Supplementary material for "A highly parallel, automated platform enabling individual or sequential ChIP of histone marks and transcription factors": SI Appendix

Table S1: primer sequences used for qPCR

| **OligoSequence(5' to 3')** | **OligoName** | **Mark tested** |
|---|---|---|
| CACACGAACCTTCCACGAG | Rplp0_TSS_F | H3K4me3 |
| TCGTTCAGCTTTGTCTGACG | Rplp0_TSS_R | H3K4me3 |
| CCACCCTGCACTTACGATG | Rplp0+200bp_F | H3K27ac |
| TGAGCTCCCTGTCTCTCCTC | RPLP0+200bp_R | H3K27ac |
| GTCCGGGTCTGACTGTCTTG | CXCL2_FW | H3K27me3, H3K9me3 |
| ACTGCACTGGGTTCACGAAG | CXCL2_RV | H3K27me3, H3K9me3 |
| GGTCATGCTGGTCTCGAACT | Rplp0-2kb_F | Negative region |
| ATCCTTCCCATGGAACACAG | Rplp0-2kb_R | Negative region |
| CGGGGGCTGCCCAAAGTTTCA | VOPP1_2_F | MEF2A, H3K4me1 |
| ATTGGGGAAATTGCAGAGCGAGC | VOPP1_2_R | MEF2A, H3K4me1 |

Supplementary Figure 1

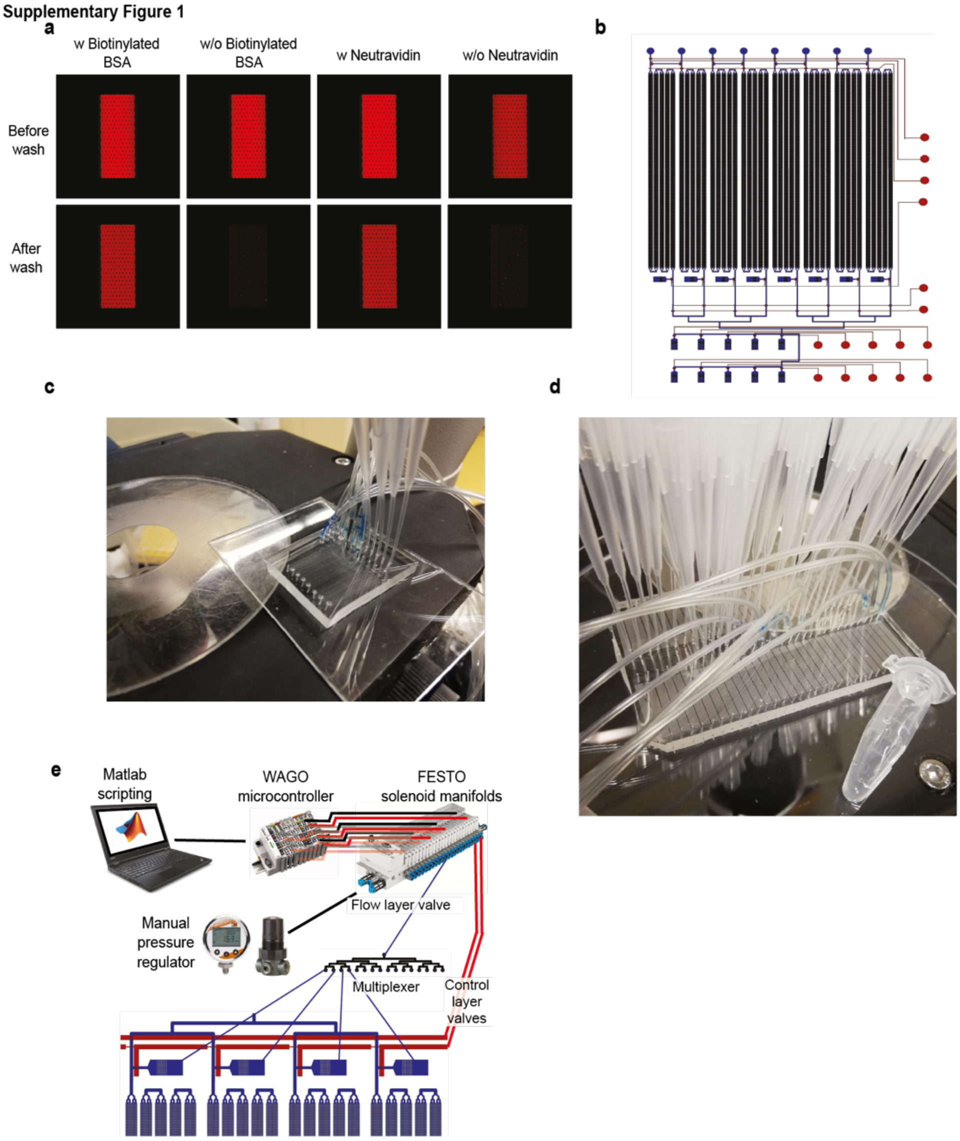


**Fig. S1**

FloChIP's setup. (**a**) Fluorescence micrographs showing the requirement for neutravidin and protein A/G in the correct formation of FloChIP's surface chemistry. (**b**) Top-view schematic of the medium-throughput 8-unit FloChIP device. Flow channels are in blue and control channels in red. (**c**) Photograph of the 8-unit device connected to pipette tips and tubing for reagent loading and pressure control, respectively. (**d**) Photograph of the 64-unit device, microcentrifuge tube for scale comparison. (**e**) Schematic of FloChIP's electronic control system.

**Supplementary Figure 2**

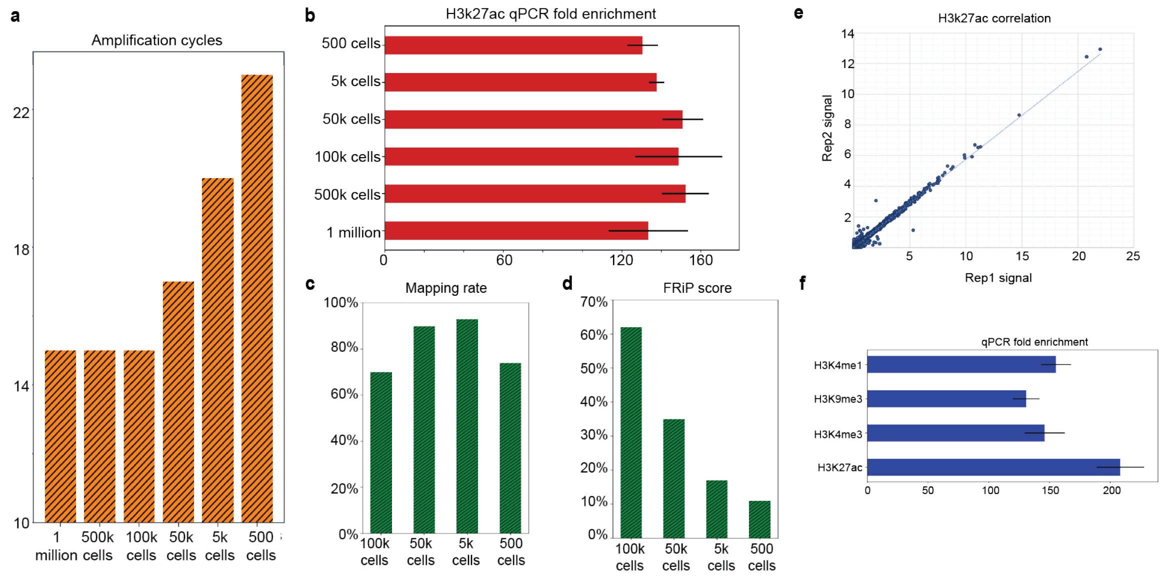


**Fig. S2**

FloChIP's results for IP on histone marks. (**a**) Number of amplification cycles required for detecting samples. Performed for several assays involving a decreasing cell number, from 1 million to 500 cells. (**b**) qPCR fold enrichment around the TSS of *RPLP0* for samples involving a decreasing cell number, from 1 million to 500 cells (error bars represent variability in qPCR replicates). (**c**) Mapping rate for samples involving a decreasing cell number, from 100k to 500 cells. (**d**) FRiP scores for samples involving a decreasing cell number, from 1 million to 500 cells. (**e**) qPCR fold enrichment for histone mark samples, namely H3K4me3 (*RPLP0* TSS), H3K27ac (*RPLP0* upstream TSS), H3K27me3 (*CXCL2* TSS), H3K9me3 (*CXCL2* TSS) and H3K4me1 (primer sequences in **Supp. Table 1**, error bars represent variability in qPCR replicates). (**f**) Correlation plot of two FloChIP libraries obtained from two FloChIP runs involving devices belonging to two distinct fabrication batches.

**Supplementary Figure 3**

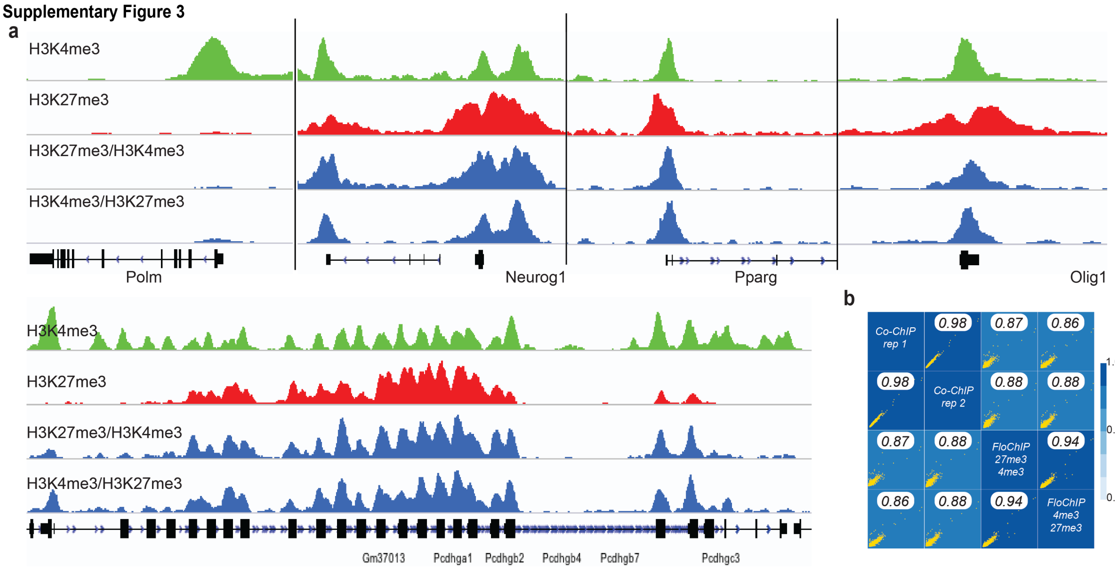


**Fig. S3**

FloChIP's sequential IP results recapitulate previously published data. (**a**) Signal tracks for individual and sequential IP libraries reported for the same five loci (four top, one bottom left) originally shown in the study by Mikkelsen and colleagues[21]. (**b**) Correlation results between FloChIP and the previously published Co-Chip data[18].

Supplementary Figure 4

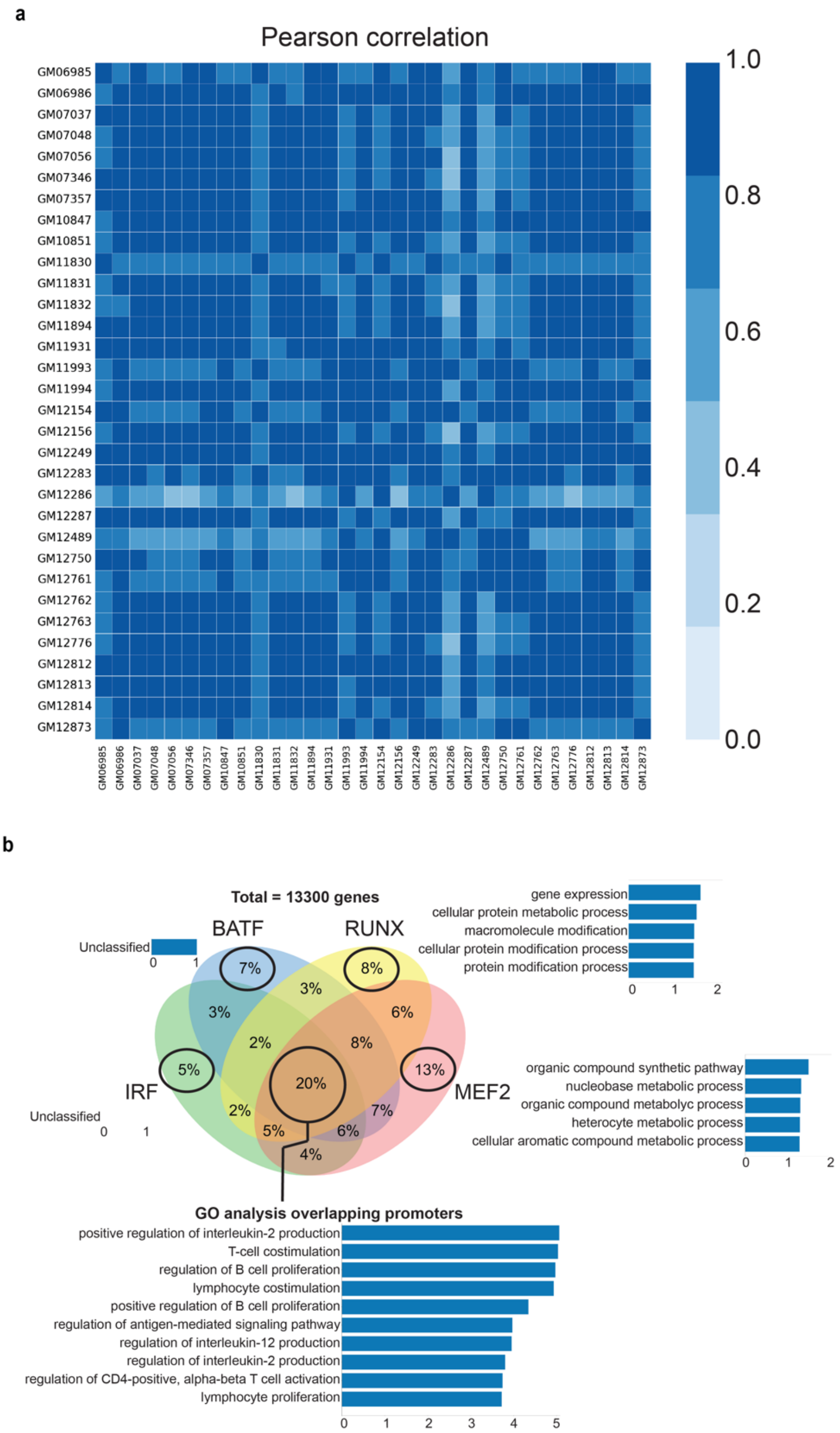


**Fig. S4**

Genome-wide analysis of FloChIP-derived MEF2A DNA binding data. (**a**) Pearson correlation between all pairs of sequenced libraries. (**b**) Gene ontology enrichment analysis of promoter sets containing individual motifs.

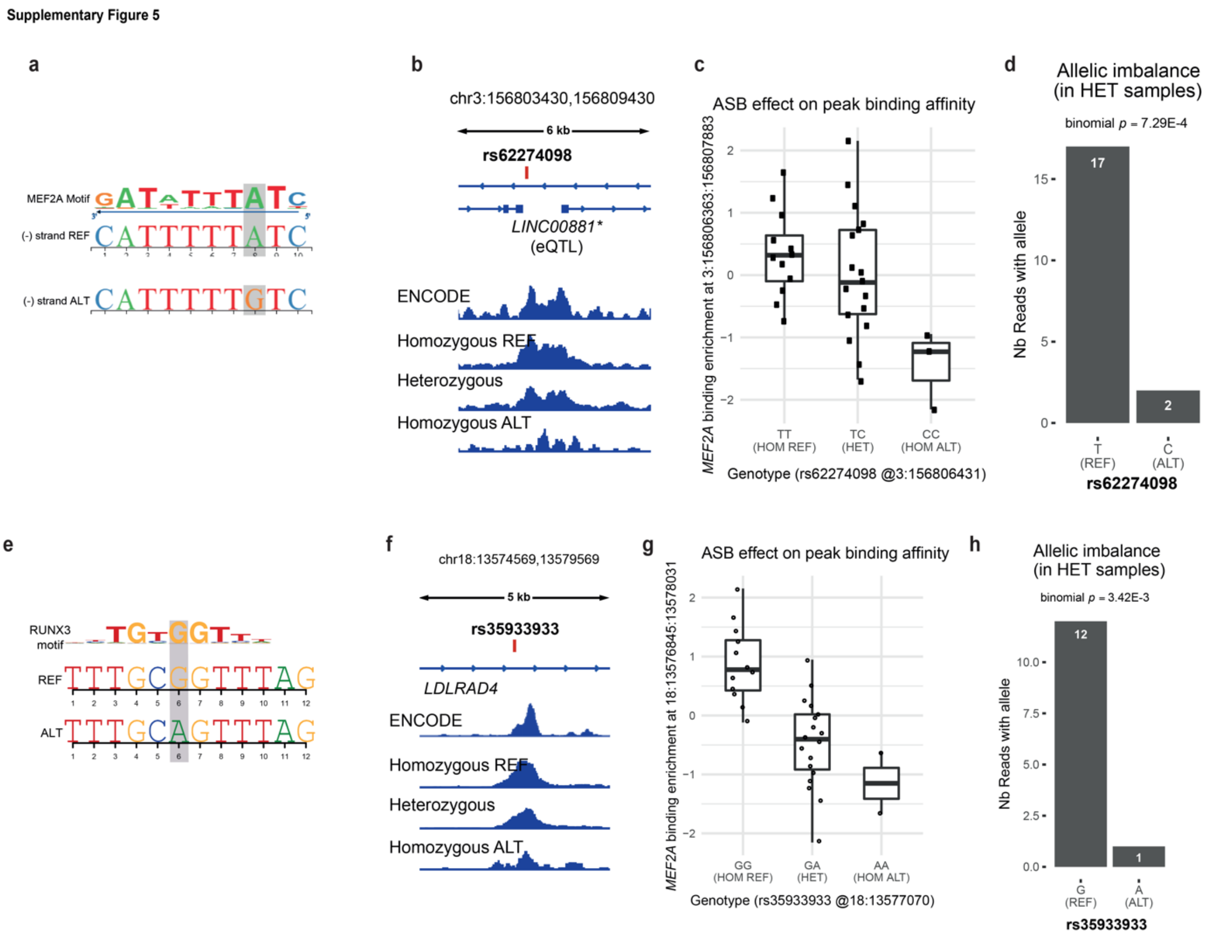


**Fig. S5**

Two MEF2A ASB examples. (**a**) rs62274098 significantly disrupts the MEF2A motif (FDR <5%). (**b**) IGV tracks of MEF2A binding enrichment at rs62274098 loci. Coverage tracks are from ENCODE (NA12878 MEF2A) and our 32 samples, stratified by rs62274098 genotype, and merged into three tracks (final bigwigs are RPKM normalized using deepTools). (**c**) Boxplots showing the effect of rs62274098 on the three different genotypes (TT, TC and CC). MEF2A binding enrichment (y-axis) is computed from the peak in which rs62274098 was found, and normalized using DESEq2, followed by qqnorm functions in R. (**d**) Allelic Imbalance highlighted for rs62274098 by summing read counts of each allele over all heterozygous samples. A binomial test yielded a p-value of 7.25E-4, revealing rs62274098 as an ASB. (**e**) rs35933933 significantly (FDR <5%) disrupts the RUNX3 motif. (**f**) IGV tracks of MEF2A binding enrichment at rs35933933 loci. Coverage tracks are from ENCODE (NA12878 MEF2A) and our 32 samples, stratified by rs35933933 genotype, and merged into three tracks (final bigwigs are RPKM normalized using deepTools). (**g**) Boxplots showing the effect of rs35933933 on the three different genotypes (GG, GA and AA). MEF2A binding enrichment (y-axis) is computed from the peak in which rs35933933 was found, and normalized using DESEq2, followed by qqnorm functions in R. (**h**) Allelic Imbalance highlighted for rs35933933 by summing read counts of each allele over all heterozygous samples. A binomial test yielded a p-value of 3.42E-3, revealing rs35933933 as an ASB.

### Supplementary methods

### Chromatin preparation

#### *Cell fixation*

Lymphoblastoid cells were harvested, washed once with PBS and resuspended in 1ml crosslinking buffer (1% PFA in PBS) for 10 minutes with shaking at RT. Crosslinking was stopped by adding 50µl of 2.5M glycine and shaking for other 5 minutes. Fixed cells were then washed twice with ice-cold PBS, pelleted, deprived of the supernatant, snap frozen and stored at -80°C.

*Lysis and sonication*

The frozen cell pellet was resuspended in ice-cold PBS at 4°C agitating for 30 minutes, spun at 1000g for 5 minutes, resuspended in lysis buffer (50 mM Hepes pH 7.8, 140 mM NaCl, 1mM EDTA, 0.5% NP40, 10% glycerol, 0.25% Triton and freshly added protease inhibitor), incubated with mild agitation for 10 minutes, spun for 5 minutes at 1000g, resuspended in nuclei wash buffer (20 mM Tris-HCl pH 8.0, 200 mM NaCl, 1 mM EDTA, 0.5 mM EGTA and freshly added protease inhibitor), incubated with mild agitation for 10 minutes, spun for 5 minutes at 1000g and resuspended in sonication buffer (20 mM Tris-HCl pH 8.0, 200 mM NaCl, 1 mM EDTA, 0.5 mM EGTA, 0.5% Na-Deoxycholate, 0.5% N-laurosylsarcosine and freshly added protease inhibitor). Nuclei were sonicated on a covaris E220 machine with the following settings: 140W intensity, 5% duty factor and 200 bursts/cycle. Chromatin was then snap frozen until ChIP.

**FloChIP**

*Device fabrication*

Microfluidic designs were generated using Tanner L-Edit and fabricated using multilayer standard soft lithography(1) at the EPFL Center for Microtechnology. Briefly, designs were first transferred to chrome masks using a VPG200 pattern generator (Heidelberg Instruments). Subsequently, microfluidic molds were assembled on silicon wafers with SU8 photoresist for the control layer and AZ9260 positive resist for the flow layer using a SUSS ACS200 Gen3 system (SUSS MicroTec). Microfluidics chips were fabricated by first separately casting PDMS onto the SU8 and the AZ9260 wafers with two different PDMS/curing agent ratios (20:1 and 5:1, respectively), partially curing for 30 minutes at 80°C, peeling off the PDMS from the AZ9260 wafer and aligning it to the SU8 wafer in order the reconstitute the desired pattern. The chips were finally fully cured at 80°C for one hour and half, peeled off, holed and plasma-bonded to clean glass slides or to PDMS-coated petri dishes.

*Experimental setup*

Automated control of the FloChIP experimental workflow is obtained by a system of components including: 1) MATLAB software, 2) a standard laptop, 3) a WAGO fieldbus controller (ModBus 750-881), 4) FESTO 3/2 way 24V miniature solenoid valves, 5) compressed air building supply (**Supp. Fig. 1e** and 6) a PCR machine. Tygon tubing and standard pipette tips are used to interface the microfluidic chip and the solenoid valves.

FloChIP is, in essence, a method consisting of the sequential introduction of different reagents into a custom-designed microfluidic chip. This sequence of reagents can be programmed with simple scripting commands that are, in turn, translated into sequences of solenoid valve actuations and releases. The concerted action of the solenoid valves, belonging to both the control layer and the flow layer of the chip, realizes in an automated fashion the required surface chemistry, immunoprecipitation and tagmentation reactions. On-chip temperature control is achieved by placing the microfluidic device on top of a PCR machine with flat heat-block and starting a pre-programmed temperature sequence in sync with the MATLAB script.

*FloChIP operation*

A FloChIP experiment starts with pre-loading the control lines with distilled water and activating all valves (at a pressure of 25-30 PSI for the control lines and 2.5-5 PSI for the flow valves). Subsequently, all the reagents required for the surface chemistry (i.e. biotin-BSA, neutravidin, PBS and biotin-protein A/G, antibodies), IP (chromatin), washes (low-salt, high-salt and LiCL buffers), tagmentation (Tn5 buffer) and elution (SDS buffer), are loaded into pipette tips and inserted into the inlets of the microfluidic device. At this stage, all valves are closed and there is no possible cross-talk between any of the reagents above. Immediately after completing the insertion of the tips, the automated protocol is launched by running the respective script. The protocol entails, in sequential order, the following steps: 20 minutes of BSA-biotin (100µl at 2mg/ml), 30 seconds of PBS wash, 20 minutes of Neutravidin (100µl at 1mg/ml), 30 seconds of PBS wash, 20 minutes of biotin-protein A/G (100µl at 2mg/ml) and 30 seconds of PBS wash.

Depending on whether direct or indirect ChIP is performed, immunoprecipitation is carried out in two different ways: direct ChIP or loading of the pre-incubated antibody/chromatin mix (indirect ChIP). For direct ChIP, following the addition of biotin-protein A/G, the antibody or antibodies of choice are loaded on chip for 20 minutes. Moreover, within direct ChIP, it is possible to operate the chip in two distinct multiplexing modes, either antibody multiplex, in which microvalves are actuated in such a way that every IP unit is functionalized with a different antibody, or sample multiplex, in which all IP units are functionalized with the same antibody (**Fig. 2a,b**). The antibodies used in this study are: Abcam antibodies: anti-H3K27ac ab4729, anti-H3K4me3 ab8580, anti-H3K4me1 ab8895, anti-H3K9me3 ab8898, anti-H3K27me3 ab6147, and anti-MEF2A sc-17785. Following antibody loading and a 1 minute PBS wash, chromatin samples are loaded on chip by opening and closing the respective microvalves. These ON/OFF cycles, usually of 2 or 5 minutes, are performed in order to ensure that the chromatin spends sufficient time inside the micropillar array for the epitopes to be efficiently recognized by the corresponding antibody.

For indirect ChIP, the antibody and chromatin are incubated for 2 or 4 hours in a PCR tube with constant mixing at 4ºC prior to loading on-chip. During the IP step, the antibody/chromatin mixes are loaded into the chip in separate IP units by utilizing the same ON/OFF cycles as mentioned above. Both for direct and indirect ChIP, the overall IP is performed at room temperature time spans between 30 and 60 minutes, depending on the amount of chromatin mix to be processed.

Following IP, rapid salt washes are performed to eliminate non-specific binding: 5 minutes of low-salt buffer (20 mM Tris pH 8.0, 150 mM NaCl, 2mM EDTA, 1% TritonX-100, 0.1% SDS), 5 minutes of high-salt buffer (20 mM Tris pH 8.0, 500 mM NaCl, 2mM EDTA, 1% TritonX-100, 0.1% SDS) and 5 minutes of LiCl buffer (20 mM Tris pH 8.0, 250 mM LiCl, 2mM EDTA, 1% NP40, 1% Na-Deoxycholate). Following these washes, Tn5 buffer (Tn5 enzyme 0.2 μM,10 mM Tris pH 8.0, 5 mM $MgCl_2$) is flown on-chip at 37°C for 45 minutes. This step ensures the complete tagmentation of the immunoprecipitated chromatin.

Following Tn5 buffer and a 5-minutes low-salt wash to remove excess adapters, SDS buffer (10 mM Tris pH 8.0, 200 mM NaCl, 1mM EDTA, 1% SDS) is loaded on-chip at 65°C for 10 minutes in order to elute the antibody-bound chromatin from the device. The eluate is independently collected from each IP lane into PCR tubes and de-crosslinked at 65°C for 4 hours. Following de-crosslinking, DNA is purified in Qiagen EB buffer using Qiagen MinElute purification kits.

*FloChIP operation for sequential ChIP*

For sequential ChIP, instead of eluting the chromatin in SDS buffer, elution is performed by saturating the antibody with a given elution peptide (ab1342 for H3K4me3 and ab1782 for H3K27me3, Abcam – Peptide elution buffer: 20µl of sonication buffer, 2µg of an antibody-specific peptide). This way, the eluted chromatin from a given IP lane can be directly re-immunoprecipitated in the subsequent IP lane. Following elution, the chromatin is collected into a pipette tip inserted in the specific chip outlet. Subsequently, by closing the microvalves connecting the first IP lane and the outlets while opening the ones connecting the outlet and the second IP lane, the chromatin is re-flown on-chip for the second IP (**Fig. 3a**) This second IP is also performed using ON/OFF cycles of 2 minutes each. The total time for the second ChIP is also between 30 and 60 minutes. Finally, after all the chromatin has been re-flown on-chip, the salt washes are repeated, tagmentation is performed and elution is achieved using the standard SDS buffer.

**ChIP-qPCR**

Following FloChIP, qPCR was used to evaluate IP efficiency prior to next generation sequencing. qPCR was performed on a StepOnePlus™ (primer sequences is Supp. Table. 1). Each qPCR reaction was composed of 10µl Applied Biosystems™ PowerUp™ SYBR™ Green Master Mix, 0.8µl of a 10µM forward primer solution, 0.8µl of a 10µM reverse primer solution, 2µl of DNA and water to a final volume of 20µl. The cycling program was the following: 2 minutes at 50°C, 2 minutes at 95°C and [5 seconds at 95°C, 20 seconds at 60°C]x60 cycles. Fold enrichment values were obtained as ratios between the percent of input of the expected positive and negative genomic regions.

**NGS Library preparation**

NGS libraries were prepared by mixing 20µl of purified DNA with 2.5µl of forward Nextera adapter, 2.5µl of reverse Nextera adapter, 32.5µl of NebNext master mix, 0.5µl of 100x SYBR green and water to 65µl. First, 5 pre-amplification cycles were run as follows: 5 minutes at 72°C, 30 seconds at 98°C and [10 seconds at 98°C, 30 seconds at 63°C, 60 seconds at 72°C]x5 cycles. Subsequently, 15µl out of the original 65µl were separated and amplified for 20 more cycles in order to estimate the optimal number of amplification cycles: 30 seconds at 98°C and [10 seconds at 98°C, 30 seconds at 63°C, 60 seconds at 72°C]x20 cycles. Finally, the remaining 50µl were amplified for N cycles, where N is the rounded up Ct value determined in the previous reaction. DNA was size selected using AMPure XP beads in order to obtain a size distribution between 150bp and 500bp. Concentrations were measured with Qubit (ThermoFisher), the size distribution was profiled with Fragment analyzer (AATI) and libraries were sequenced on an Illumina NextSeq 500.

**FloChIP read mapping and processing for histone marks and TFs**

Sequencing reads were mapped to the human (hg38) and mouse (mm10) genomes using STAR(2) with default parameters. Uniquely mapped reads were used to call peaks using the HOMER(3) command *findPeaks.pl* with the appropriate flag, i.e. *–histone* for histone marks and *–factor* for transcription factors. FRiP scores were calculated using HOMER's command *annotatePeaks.pl*, dividing the total number of reads that fall within peaks by the total number of mapped reads. Correlation plots were generated using *annotatePeaks.pl* on a common peak file, either ENCODE's peak files or, alternatively, the overlapping set of peaks between ENCODE and FloChIP datasets. For ChIP-seq on TFs, reads were mapped to the human genome assembly hg19 – for consistency with previous ChIP-seq studies on the same cell lines(4). TF peak calling was performed with HOMER's script *findPeaks.pl* with default parameters and the flag *-factor.* The *de novo* and known motif enrichment results were obtained with the HOMER's function *findMotifsGenome.pl*, executed for each peak file with default parameters. The percentage of peaks containing specific motifs was found by using the function *annotatePeaks.pl* with the option -m followed by the given motif file.

**Bivalency score calculation**

The bvScore is assigned to each promoter and is intended to consider both the intersection between two ChIP-seq datasets as well as the agreement between the respective sequential ChIP-seq datasets. Accordingly, the bvScore can be expressed as the product of the co-occurrence score (cScore), which measures the relative coverage of the two ChIP-seq tracks, and the agreement score (aScore), which measures the relative coverage of the two sequential ChIP-seq tracks. We define the cScore as $(nmr_{4i}+nmr_{27i})/(abs(nmr_{4i}-nmr_{27i})+1)$, where $nmr_{4i}$ and $nmr_{27i}$ are the normalized number of mapped reads in promoter *i* for H3K4me3 and H3K27me3, respectively. The higher the value of the cScore for a promoter, the more similar is the occupancy of the two marks on that promoter. We define the aScore as the absolute value of $(nmr_{4/27i}+nmr_{27/4i})/(abs(nmr_{4/27i}-nmr_{27/4i})+1)$, where $nmr_{4/27i}$ and $nmr_{27/4i}$ are the number of mapped reads on promoter *i* for the two sequential ChIP-seq experiments. The higher the aScore of a promoter, the more similar is the coverage of the two sequential ChIP-seq datasets on that promoter. Finally, the bivalency score is thus defined as $bvScore=log(abs(cScore*aScore))$.

Gene ontology analysis was performed using the online tool http://geneontology.org/page/go-enrichment-analysis.

**Allele-Specific Binding**

For identifying variants subject to Allele-Specific Binding of MEF2A (ASB), we first preprocessed the BAM files according to GATK's Best Practices recommendations[29] including duplicate removal using Picard v.2.17.8 (http://broadinstitute.github.io/picard/). We downloaded the 1000G genotyping data from the EBI server (hg19), removed variants with a minor allele frequency lower than 5%, while restricting our analysis to the 32 samples of interest, which yielded a total of 1,268,985 single nucleotide polymorphisms (SNPs). Then, we applied the *ASEReadCounter* tool from GATK v.4.0.4.0, on each of the 32 samples. These results were then merged by summing read counts for every SNP across heterozygous samples. We filtered out all SNPs with a coverage lower than 10 reads, which yielded 6,330 SNPs. Then we opted to keep only SNPs falling into called peaks, which led to 4,554 SNPs that were further analyzed. On these, we performed a binomial test to assess significant allelic imbalance (nominal p-value 5%), yielding 751 potential ASBs (37 at FDR 5%).

**TF motif disruption analysis**

All downstream analyses were performed using R v. 3.5.0. We analyzed all 751 ASBs to check if they were significantly impacting Transcription Factor (TF) motifs using atSNPv.1.0(5) with the *BSgenome.Hsapiens.UCSC.hg19* package as genome library and *SNPlocs.Hsapiens.dbSNP144.GRCh37* package as SNP library. We tested 401 mononucleotide human core TF motifs that were downloaded from HOCOMOCO v11(6).

**Allele Binding Cooperativity**

Allele Binding Cooperativity (ABC) was assessed for all 751 ASBs using a linear regression between the motif disruption/creation log likelihood ratio computed by atSNP (see previous Methods section) and the fold change between the ASB ref and alt counts. This analysis was performed for all SNPs significantly disrupting any of the 401 motifs analyzed by atSNP at FDR5%, and allowed to find which ASBs were concordant with motif disruption across several tested SNPs.
